# Development of an ELISA-Based Method for Testing Aflatoxigenicity and Aflatoxigenic Variability among *Aspergillus* species in Culture

**DOI:** 10.1101/523506

**Authors:** Lagat Kipkemboi Micah, Faith Jebet Toroitich, Meshack Amos Obonyo

**Author notes:** These authors contributed equally to this work.

## Abstract

Aflatoxins contaminate foodstuff posing a severe threat to human health because chronic exposure is linked to liver cancer while acute exposure may cause death. Therefore, it is of interest to reduce the contamination of crops by aflatoxins in the field and post-harvest. Among the current technologies being developed is the deployment of non-aflatoxigenic strains of *Aspergillus* species to competitively exclude aflatoxigenic conspecifics from crops in the field thereby curtailing aflatoxin production by the former. The success in this endeavor makes the non-aflatoxigenic fungi good candidates for biological control programs. However, the current techniques for segregating non-aflatoxigenic from aflatoxigenic fungi suffer two main drawbacks: they are based on morphological and chemical tests with a combination of visual color changes detected in a culture plate which suffer some degree of inaccuracy. Secondly, the existing methods are incapable of accurately quantifying aflatoxin production by fungi in culture. We developed a culture system for inducing aflatoxin production by *Aspergillus* using maize kernels as growth substrate followed by quantification using ELISA. The method was compared to the Dichlorvos-Ammonia (DV-AM) method for determining aflatoxigenicity. Our findings encapsulate a method more robust than the currently used DV-AM approach because, for the first time, we are able to assess aflatoxigenicity and aflatoxigenic variability among *Aspergillus* species earlier classified as non-aflatoxigenic by the DV-AM method. Furthermore, the new method presents an opportunity to attribute toxin production by actively growing fungal cultures. We believe this method when further developed presents a chance to study and predict fungal behavior prior to field trials for biological control programs.

## 1.0 Introduction

Aflatoxin contamination of foodstuff continues to pose a serious risk to human health when contaminated food material gets into the food chain and is eventually consumed by humans [1]. Apart from the health concern, the contamination of food material by aflatoxins presents a significant barrier to trade. This is because the contaminated commodity is rejected in the export market leading to losses of revenue currently estimated at US$ 670 Million in Africa alone [2]. There are several management options that have been proposed and tested to mitigate aflatoxin contamination of crops both in the field and after harvest. One of the most promising strategies undergoing development is use of non-aflatoxin producing (non-aflatoxigenic) fungi species in the genus *Aspergillus* to control aflatoxin producing (aflatoxigenic) relatives in the field [3].

In order to be successful, such biological control program based on fungi-fungi interaction must of essence first, correctly identify candidate non-aflatoxigenic strains of *Aspergillus* species; then accurately predict their behavior while interacting with aflatoxigenic fungi in field conditions. To identify candidate fungi for use in biological control formulations, researchers rely on diverse morphological, chemical [4], and to some extent a combination of molecular and metabolic methods to establish aflatoxigenicity [5]. The aforementioned methods currently in use enjoy some degree of success as well as inconsistencies in reporting non-aflatoxigenic fungi thus leaving a knowledge gap occasioned by the inconclusive nature of their findings.

To the best of our knowledge, since the introduction of the suspended disc method by Norton [6], there has been no other technique developed to quantify aflatoxin production by viable and growing fungi cultures. The widely accepted method has been the use of conventional (YES) growth media as a substrate to induce aflatoxin production [4, 7]. Followed by, in the alternative, the Dichlorvos-Ammonia (DV-AM); a chemical-visual method of detection of aflatoxigenic fungi recently improved by Kushiro, Hatabayashi [8].

In an effort to advance the existing techniques, we present a robust method based on its ability to utilize natural growth substrate to induce and quantify aflatoxin production and in tandem give information on aflatoxigenicity and aflatoxigenic variability among viable and actively growing *Aspergillus* cultures.

## 1.2 Materials and Methods

### 1.2.1 Culture of fungi

The *Aspergillus* species used in this study had previously been isolated from maize growing in Eastern Kenya [9]. The tested fungi were retrieved from storage and sub-cultured on Potato Dextrose Agar (PDA) media (HiMedia Laboratories Pvt. Ltd). The PDA media was amended with 50mg/L streptomycin sulfate and penicillin (Zhonghuo Pharmaceuticals, China) to inhibit bacterial growth. Inoculation was done, and fungi were grown in Petri dishes (90 × 15mm; Aptaca™, Italy) containing the growth media. The set up was incubated at 28°C for seven days. Dichlorvos (Amiran Kenya Ltd) was used to set up the DV-AM method. For the DV-AM method, the retrieved samples were cultured on aflatoxin inducing YES media as described by Kushiro, Hatabayashi [8].

### 1.2.2 Preparation of maize kernel growth substrate

Maize kernels without any signs of damage were used as a substrate for fungal growth. The kernels were first surface sterilized in 25 % (v/v) sodium hypochlorite (NaOCl) for 15 minutes then rinsed in sterile distilled water. Subsequently, the kernels were treated with ammonium hydroxide (NH_4_OH) for 3 hours (to reduce and standardize endogenous aflatoxins in the growth substrate making it suitable as a blank material), the process adopted from Namazi, Allameh [10]. Afterward, the kernels were rinsed in sterile distilled water three times. The maize kernels were then introduced into a laminar flow overnight under ultraviolet light to prevent possible fungal contamination (this ensured only the introduced fungi grow). Afterward, approximately 25 grams of the sterile maize kernels were introduced into sterile 25 mL sample bottles (Rudolph Research Analytical).

### 1.2.3 Induction of aflatoxin production by fungi cultures

The PDA culture plates with growing fungi were irrigated with 3 mL of sterile (autoclaved) distilled water and gently swirled to dislodge conidia from the surface of the colonies. 1 mL of water was carefully aspirated with a pipette, and the spores counted using a Petroff-Hausser cell-counting chamber (Hausser Scientific). The conidia concentration was then adjusted to 1× 10^6^ mL^−1^; after which 500 μL of the water with the conidia suspension was sprinkled into the maize kernel packed bottles and covered ready for incubation. A negative control consisting of treated maize samples with 500μL of sterile water without fungal conidia was set up [Fig 2 (B)]. The samples and controls were incubated at 28°C for seven days. During the incubation period, 100 μL of sterile water was sprinkled at two-day interval into the packed maize bottles to prevent desiccation. The experiment was replicated thrice.

### 1.2.4 Quantification of aflatoxin production

On the 8th day (after the seven-day incubation period), samples were withdrawn and dried in an oven for 24 hours at 40°C and then pulverized into a fine powder using a manual hand-grinder (Whatman® Inc.). For quantification, the competitive ELISA procedure for quantification of total Aflatoxins was employed. Briefly, 20g of the maize powder was mixed with 100mL methanol (70% v/v) to extract aflatoxins. The mixture was filtered (Whatman #1) from which 10mL of the filtrate aliquoted into 14mL BD Falcon™ test tubes (BD Biosciences, Bedford, USA) and labeled accordingly. Total Aflatoxins were analyzed using ELISA kit (Helica Biosystems Inc®, Santa Ana-USA) following manufacturer’s instructions and absorbance read at 450nm in a micro-plate reader (Mindray® Inc. Nanshan, Shenzhen, China).

### 1.2.5 Analyses

A one-sample t-test was performed to compare the aflatoxin levels between samples and controls. The statistical analysis was carried out using the Statistical Package for Social Science (SPSS) ver. 20.0 (IBM Corp., Armonk, NY). All the data with *t* (6)>1.943, *p*< 0.05 were considered significant. The images of the sample plates and sample bottles with fungal growth were analyzed using ImageJ 1.x software [11].

### 1.2.6 Molecular methods

Fungi Isolates to be subjected to molecular identification were grown in Potato Dextrose Agar (PDA; pH 6.0). The isolates were maintained at 28°C for five days. The periphery of the exponentially growing fungal mycelium (50-100 mg) was excised aseptically using a sterile scalpel. DNA extraction was done using the Quick-DNA™ Fungal/Bacterial Kit (Zymo Research). Finally, 100 μL of DNA was eluted for DNA amplification. Specific primers for *aflQ* (Forward- (5′-TTA AGG CAG CGG AAT ACA AG-3′) and Reverse- (5′-GAC GCC CAA AGC CGA ACA CAA A-3′)) were synthesized by Inqaba Biotechnical Industries (Pty) Ltd (South Africa).

A total reaction volume of 25μL consisting of 12.5μL of One Taq^®^ 2X Master Mix (New England Biolabs), 0.2 μL of DNA template (< 1000 ng), 0.5 μL (0.2 μM) of each (forward and reverse) of the primers and 9.5 μL of nuclease-free water. The mixture was gently mixed by priming pipettor at least 4 times and spinned prior to PCR. PCR tubes were transferred to a preheated thermocycler at 94 °C for 3 minutes followed 30 cycles consisting of 1 minute of denaturation at 94°C, 1 minute of annealing at 57°C and 1 minute of extension at 72°C. The final extension of 10 minutes at 72°C and the storage temperature of 4°C were considered. The PCR products were subjected to gel electrophoresis using 1% agarose gel pre-cast with ethidium bromide. The optimized conditions for the process involved a constant supply of voltage (50 V (41mA)) for 75 minutes in a tris-borate-EDTA (TBE) buffer electrophoretic chamber. The gel was visualized using Automatic Gel Imaging System (Peiqing Science and Technology Co., Ltd).

## 1.3 Results

In general, there was positive fungal growth in the sample bottles (maize culture system) as well as the culture plates during the incubation period [Fig 1]. There was aflatoxin production by the fungi incubated in the maize culture system (ELISA analyzed) as well as the YES media culture plates (Image J-analyzed) [Fig 3]. Upon visualization of the YES media culture, plates using the DV-AM method, three fungi isolates were classified as non-aflatoxigenic while another four were aflatoxigenic. By contrast, and to our surprise, all fungi were recorded as aflatoxigenic in the maize culture system when the substrate was tested in ELISA [Table 1]. In general, there was an increase in the amount of aflatoxins in the samples compared to the treated control implying variation in aflatoxigenicity of the fungi. Two of the *Aspergillus flavus* (1EM1901 and 1EM1201) isolates produced significantly high aflatoxin levels above the regulatory limit (of 10 ppb) implying aggressive production of aflatoxins whereas the DV-AM method showed that the isolate (1EM1201) was non-aflatoxigenic due to lack of the visible red coloration on the reverse of the culture plate. Another intriguing observation based on the aflatoxin producing capability was observed with the presence of aflatoxin-associated gene (*aflQ* (*ordA*)) across all the isolates [Fig 4].

**Fig 1.**
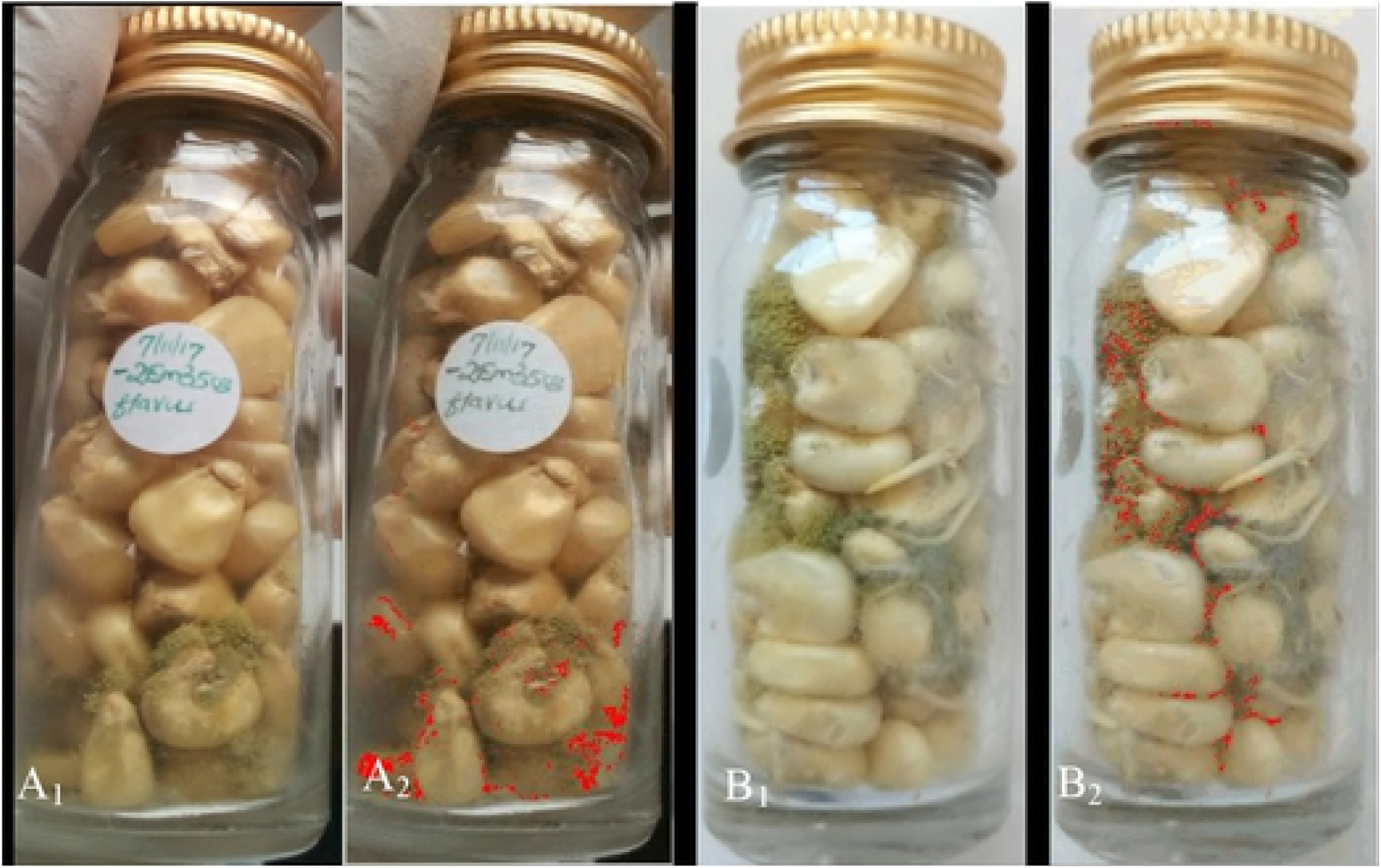
Seven-day-old fungi isolates inside the bottles before (A_1_ and B_1_) and after (A_2_and B_2_) ImageJ 1 .x processing; Red highlights in (A_2_ and B_2_) show the regions of fungal growth;

**Fig 2.**
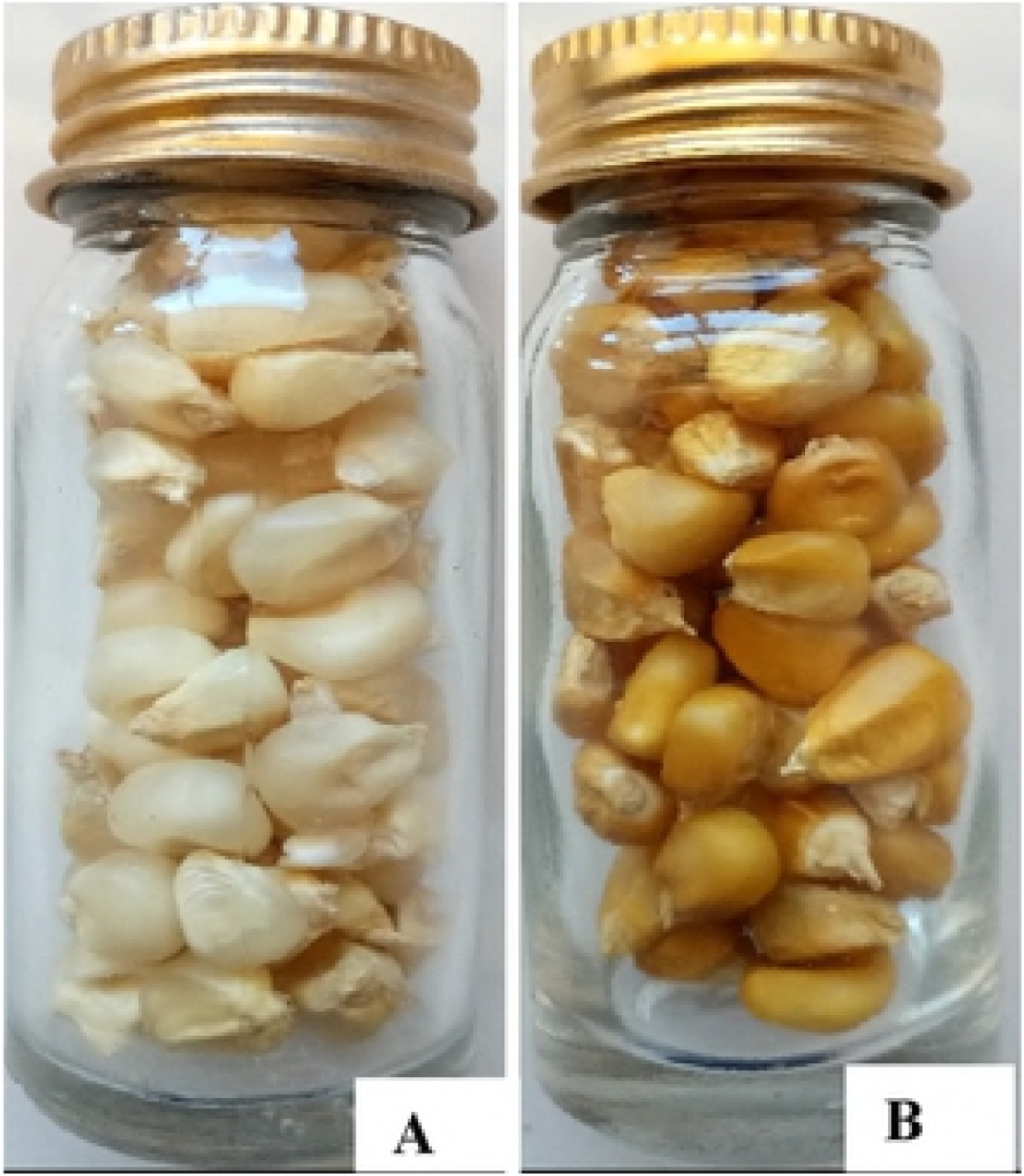
Set up of the Maize Kernel culture system (A) untreated and (B) ammonia-treated maize.

## 1.4 Discussions

The ELISA-based maize culture system herein reported has for the first time demonstrated aflatoxigenic variability among congeneric species of *Aspergillus* thereby separating the fungi on their aflatoxigenic ability. This is made possible by use of the maize substrate that mimics natural contamination enabling downstream quantification using ELISA. Furthermore, this method ensures the viability of maize kernels is maintained as they germinated at the end of the experiment. The current results have demonstrated that there are some amount of aflatoxins produced by fungi in a culture that may not be reported using conventional DV-AM methods. The presence of aflatoxin-associated gene (*aflQ* (*ordA*)) confers the inherent capabilities of the fungi isolates to produce aflatoxins just as in the case of the newly developed method. The (*aflQ* (*ordA*)) gene encodes for an oxidoreductase enzyme that converts O-methylsterigmatocystin (OMST) and dihydro-O-methylsterigmatocystin (DHOMST) to aflatoxins (B1, B2, G1, and G2) as described by Yu, Chang [12]. The gene is an excellent indicator that the intemediates of aflatoxins biosynthesis are ultimately converted to aflatoxins.

Despite the levels of aflatoxins being statistically insignificant in some cases, the increase when compared to the treated control is considered substantial. This finding, we believe, is an important milestone to be considered in the design of biological control programs [13], that make use of non-aflatoxigenic fungi species to competitively exclude aflatoxigenic conspecifics thereby reducing aflatoxin contamination in the field [14].

An important observation that deserves mention is that *Aspergillus* species already considered non-aflatoxigenic using the DV-AM method; when incubated in the novel maize culture system produced detectable amounts of aflatoxins. To the best of our knowledge, there has not been a simpler, cost-effective and sensitive quantitative method of estimating aflatoxin production by viable cultures of *Aspergillus* species. This information is important considering the phenomenal growth of biological control programs targeting the reduction of aflatoxin contamination in food material not only in Africa but across the globe. However, to date, the study of aflatoxigenicity and aflatoxigenic variability among *Aspergillus* species relies on morphological [13] and chemical characteristics [8] as well as a complex combination of metabolic and molecular methods of assessment [5, 8]. To some extent, these different methods have returned an inconclusive verdict on aflatoxigenic variability and aflatoxigenicity among *Aspergillus* species. Additionally, there has not been a method that yields information on aflatoxigenic variability and aflatoxigenicity concurrently in a single set-up. An important knowledge gap has been to design an experiment that closely mimics natural events in the field especially given the very high fungal diversity in the field Islam, Callicott [15].

## 1.5 Conclusion

The ELISA-based maize culture system provides a simpler and sensitive method for inducing aflatoxin production *in vitro* among fungal isolates that could otherwise remain undetected using DV-AM method. It offers the best model to simulate field interactions among fungi. In view of the fact that DV is a highly toxic and regulated chemical, the current method suffices to study aflatoxigenic variability and aflatoxigenicity among *Aspergillus* species both in resource-limited and limitless settings. While ELISA-systems may suffer shortcomings of reporting false positives, the quantitative aspect can be used in HPLC system for more accurate estimation on toxins.

## 1.7 Authors Contributions

LKM performed the experiments, generated data, and drafted the manuscript; MAO and FJT designed the research study, sourced funding, supervised the project and reviewed the manuscript. All authors approved the final manuscript for submission.

## Acknowledgments

The experiments were conducted at the Mycotxin Research Laboratory at Egerton University supported by the Canadian Government through the Grand Challenges Canada (Grant ID S7 0656-01-10)

## References

1. Obonyo MA, Salano EN. Perennial and seasonal contamination of maize by aflatoxins in eastern Kenya. International Journal of Food Contamination. 2018;5(1):6.

2. PACA. First progress report of the chairperson of the commission on food safety. 2018.

3. Damann Jr KE. Atoxigenic *Aspergillus flavus* biological control of aflatoxin contamination: what is the mechanism? World Mycotoxin Journal. 2015;8(2):235–44.

4. Rodrigues P, Venâncio A, Kozakiewicz Z, Lima N. A polyphasic approach to the identification of aflatoxigenic and non-aflatoxigenic strains of *Aspergillus* section *Flavi* isolated from Portuguese almonds. International journal of food microbiology. 2009;129(2):187–93.

5. Okoth S, De Boevre M, Vidal A, Diana Di Mavungu J, Landschoot S, Kyallo M, et al. Genetic and Toxigenic Variability within *Aspergillus flavus* Population Isolated from Maize in Two Diverse Environments in Kenya. Frontiers in Microbiology. 2018;9(57).

6. Norton RA. A novel glass fiber disc culture system for testing of small amounts of compounds on growth and aflatoxin production by *Aspergillus flavus*. Mycopathologia. 1995;129(2):103–9.

7. Okoth S, Nyongesa B, Ayugi V, Kang’ethe E, Korhonen H, Joutsjoki V. Toxigenic Potential of *Aspergillus* Species Occurring on Maize Kernels from Two Agro-Ecological Zones in Kenya. Toxins. 2012;4(12):991–1007.

8. Kushiro M, Hatabayashi H, Yabe K, Loladze A. Detection of Aflatoxigenic and Atoxigenic Mexican *Aspergillus* Strains by the Dichlorvos–Ammonia (DV–AM) Method. Toxins. 2018;10(7):263.

9. Salano EN, Obonyo MA, Toroitich FJ, Odhiambo B, Omondi, Aman B, Omondi. Diversity of putatively toxigenic *Aspergillus* species in maize and soil samples in an aflatoxicosis hotspot in Eastern Kenya. African Journal of Microbiology Research. 2016;10(6):172–84.

10. Namazi M, Allameh A, Aminshahidi M, Nohee A, Malekzadeh F. Inhibitory effects of ammonia solution on growth and aflatoxins production by *Aspergillus parasiticus* NRRL-2999. Acta Poloniae Toxicologica. 2002;1(10).

11. Schneider CA, Rasband WS, Eliceiri KW. NIH Image to ImageJ: 25 years of image analysis. Nature methods. 2012;9(7):671.

12. Yu J, Chang P-K, Ehrlich KC, Cary JW, Montalbano B, Dyer JM, et al. Characterization of the Critical Amino Acids of an *Aspergillus parasiticus* Cytochrome P-450 Monooxygenase Encoded by *ordA* That Is Involved in the Biosynthesis of Aflatoxins B1, G1, B2, and G2. Applied and environmental microbiology. 1998;64(12):4834–41.

13. Cotty PJ, Antilla L, Wakelyn PJ. Competitive exclusion of aflatoxin producers: farmer driven research and development. Biological control: a global perspective. 2007:241–53.

14. Damann K. Atoxigenic *Aspergillus flavus* biological control of aflatoxin contamination: what is the mechanism? World Mycotoxin Journal. 2014;8(2):235–44.

15. Islam M-S, Callicott KA, Mutegi C, Bandyopadhyay R, Cotty PJ. *Aspergillus flavus* resident in Kenya: High genetic diversity in an ancient population primarily shaped by clonal reproduction and mutation-driven evolution. Fungal Ecology. 2018;35:20–33.

